# Feeding status influences adult vine weevil (coleoptera: curculionidae) behaviour to conspecific volatiles

**DOI:** 10.1101/789172

**Authors:** Joe M. Roberts, Jhaman Kundun, Tom W. Pope

## Abstract

Vine weevil, *Otiorhynchus sulcatus* F. (Coleoptera: Curculionidae), is widely considered to be one of the most economically important pests of soft-fruit and ornamental crops. The chemical ecology of vine weevil has been extensively studied in the pursuit of a semiochemical lure to improve monitoring tool sensitivity/reliability and develop novel control methods. Although vine weevil adults exhibit a strong tendency to aggregate, the mechanism underlying this behaviour has not, to date, been identified. It is notable, however, that previous studies have not considered the impact of feeding status on vine weevil aggregation behaviour. To investigate the importance of feeding status on aggregation behavior, this study recorded the responses of satiated and starved vine weevil adults to the odour of starved conspecifics. Satiated adults exhibited a preference for the odour of conspecifics while starved vine weevils exhibited no such preference. Therefore, this study provides evidence to support the hypothesis that feeding status is an important factor for vine weevil aggregation.

## INTRODUCTION

Vine weevil, *Otiorhynchus sulcatus* F. (Coleoptera: Curculionidae), is one of the most economically important pests of soft-fruit and ornamental crops (van Tol, Bruck, Griepink & De Kogel, 2012). Adults are flightless and deposit their eggs into cracks in the soil or growing medium at night (Smith, 1932), where, upon hatching, the larvae complete four to nine molts prior to pupation (Masaki & Ohto, 1995). Crop damage and economic losses are the result of feeding on plant roots, corns and rhizomes by larvae and on leaves by adults (Moorhouse, Charnley & Gillespie, 1992). Control of vine weevil adults has typically relied on foliar applications of pyrethroid insecticides while control of larval stages has been through incorporation of slow release granules of neonicotinoid insecticides (Karley, Shepherd, Hall, McLaren & Johnson, 2012). However, larval control has now shifted in many situations toward replacing these insecticides with entomopathogenic nematodes and fungi with some success (Ansari, Shah & Butt, 2008).

The chemical ecology of vine weevil has been extensively studied in the pursuit of a semiochemical lure to improve monitoring tool sensitivity/reliability and develop novel control methods, such as autodissemination of entomopathogenic fungi (Pope et al., 2018). As vine weevil reproduce via thelytokous parthenogenesis (Lundmark, 2010), and are subsequently all female, they do not produce a sex pheromone (van Tol et al., 2012). Although vine weevil does not produce a sex pheromone, it is thought an aggregation pheromone may exist as adult vine weevils exhibit aggregation behaviour and a preference for volatiles originating from conspecifics in olfactometer experiments (Pickett, Bartlett, Buxton, Wadhams & Woodcock, 1996; Kakizaki 2001; Tol et al., 2004; Nakamuta et al., 2005). However, studies so far completed focus only on the behavioural response of starved individuals. To address this gap in our understanding of vine weevil behavior we investigated the effect of feeding status on the behavioural responses of vine weevil adults to the odour produced by conspecifics.

## MATERIALS AND METHODS

### Insect cultures

Adult vine weevils, *Otiohynchus sulcatus* F., were collected from commercial strawberry crops grown in Shropshire and Staffordshire (UK) during the summer in 2017. Recovered vine weevils were maintained on branches of yew, *Taxus baccata* (L.), and moist paper towels inside insect cages (47.5 × 47.5 × 47.5 cm) (Bugdorm, MegaView, Taiwan) placed in a controlled environment room (20 °C; 60% RH; L:D 16:8 h) (Fitotron, Weiss Technik, Ebbw Vale, Wales).

### Y-tube olfactometer bioassays

Behavioural responses of starved and satiated adult vine weevils were recorded to the volatiles from 40 starved conspecifics that had been starved for 24 hours prior to the start of the experiment. Air passing through the olfactometer was charcoal filtered air. The Y-tube olfactometer was made of glass (Sci-Glass Consultancy, UK) based upon the design used by van Tol, Visser & Sabelis (2002) with a stem length of 120 mm, arm length 190 mm and an internal diameter 18 mm. Prior to use the olfactometer was ‘smoke-tested’ for airflow visualisation to ensure odour fields were discrete. The airflow in each arm was 600 ml/min and conspecifics acting as an odour source were held in a 500 ml Dreschel bottle (Sci-Glass Consultancy, UK) for at least two hours before an experiment began, an empty Dreschel bottle was used for the control.

All bioassays were done in darkness under controlled environmental conditions (20 °C; 60 % RH; 16:8 h L:D). Prior to use in a bioassay, vine weevils were either starved for a minimum of 24 hours or collected from their culture cages and used immediately. Groups of 40 vine weevils (starved or satiated) were introduced into the olfactometer via a release chamber (100 mm diameter; Sci-Glass Consultancy, UK). The experiment was replicated four times with fresh individuals. Each replicate lasted for a maximum of 20 min and the number of weevils reaching the end of each arm during this time was recorded. Odour source positions were alternated between replicates to eliminate directional bias. After each pair of odour sources had been tested glassware was cleaned by rinsing with warm water followed by HPLC-grade acetone (Sigma Aldrich, UK), then baked in a glassware oven at 120 °C for 15 min.

### Statistical analyses

All statistical analyses were performed using R (Version 3.5-3; R Core Team, 2019). Y-tube olfactometer bioassay data was analysed using an exact binomial test against the null hypothesis that the number of vine weevils reaching the end of either olfactometer arm had a 50:50 distribution. Prior to statistical analysis, replicated results from each feeding status tested were pooled and non-responding individuals were excluded.

## RESULTS

Satiated vine weevil adults exhibited a preference for the Y-tube olfactometer arm containing air blown over 40 starved conspecifics with 74 % of responding individuals choosing this arm over the clean air control arm (*P* < 0.001) while starved adults exhibited no such preference with 44 % choosing the arm containing conspecific volatiles (Fig. 1).

**Figure 1.**
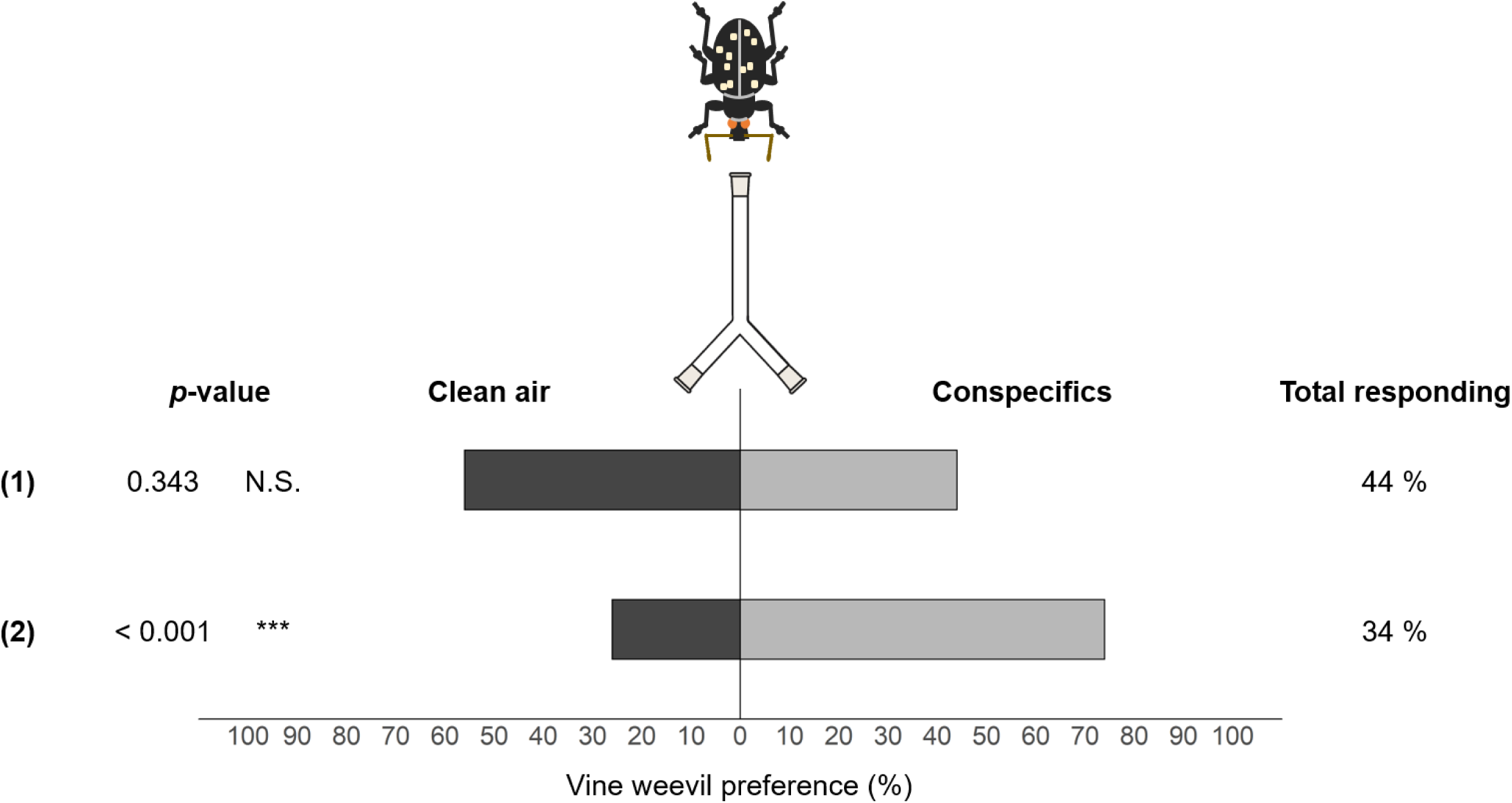
Behavioural response of adult vine weevil when presented with a choice between two odours (*n* = 160). The feeding status of the vine weevils being presented with the choice are: (1) starved and (2) satiated. Asterisks indicate a preference that is significantly different from the expected distribution of 50:50 using a binomial exact test: *** *P* < 0.001; N.S. = not significant.

## DISCUSSION

During daylight hours, vine weevil adults are often found aggregating within refuges close to their host plants. Understanding vine weevil aggregation has been the subject of extensive research, with results suggesting that this behaviour is likely to be mediated by volatile cues originating either from the frass produced by conspecifics (Pickett et al., 1996; van Tol, Visser & Sabelis, 2004) or the conspecifics themselves (Kakizaki 2001; Nakamuta, van Tol & Visser, 2005). Previous work, however, has largely focussed on the behavioural response of starved weevils toward conspecific volatiles, which is likely to be an unrealistic situation as vine weevil adults aggregate at dawn after feeding (Moorhouse et al., 1992).

It is possible that any positive behavioural response to the frass produced by conspecifics by starved vine weevils is a response toward plant volatiles from undigested plant material within the frass rather than to the weevils themselves. Vine weevil frass is known to contain volatile organic compounds (VOCs) commonly emitted by plants (e.g. *β*-caryophyllene, linalool, eugenol, etc.) (Karley et al., 2012), many of which have been implicated in host-plant selection in previous studies (Karley et al., 2012; van Tol et al., 2012). In the study presented here, conspecifics used as an odour source were starved for 24 hours prior to the start of the experiment to avoid the production of frass. In this way satiated weevils responding to the odour of conspecifics were doing so in response to volatile cues produced directly from the weevils rather than a mixture of cues produced by weevils and their frass. As starved vine weevils showed no preference for conspecific volatiles in this study, it seems likely that the positive behavioural responses of starved weevils to conspecifics reported in previous studies was due to the presence of frass volatiles (Pickett et al., 1996; Tol et al., 2004). As satiated vine weevil adults exhibited a preference for conspecific volatiles in the absence of frass in this study there is, for the first time, evidence of a positive behavioural response to volatiles produced directly by conspecifics.

This study highlights the importance of feeding status to vine weevil behavioural responses and provides new information that may help inform development of new monitoring tools for management strategies. Enhancing our understanding of how feeding status influences aggregation could lead to the identification of an aggregation pheromone by providing optimum experimental conditions for pheromone collection and analysis.

## ACKNOWLEDGEMENTS

The authors thank Dr Andrew Beacham for help in preparing Figure 1. This work was funded by AHDB Horticulture [Project number HNS 195].

## AUTHOR CONTRIBUTION

- JMR and TWP conceived research.
- JK conducted experiments.
- JMR analysed data and conducted statistical analyses.
- JMR, JK and TWP wrote the manuscript.
- TWP secured funding.
- All authors read and approved the manuscript.

